# High throughput, high fidelity genotyping and *de novo* discovery of allelic variants at the self-incompatibility locus in natural populations of Brassicaceae from short read sequencing data

**DOI:** 10.1101/752717

**Authors:** Mathieu Genete, Vincent Castric, Xavier Vekemans

## Abstract

Plant self-incompatibility (SI) is a genetic system that prevents selfing and enforces outcrossing. Because of strong balancing selection, the genes encoding SI are predicted to maintain extraordinary high levels of polymorphism, both in terms of the number of S-alleles that segregate in SI species and in terms of nucleotide sequence divergence among distinct S-allelic lines. However, because of these two combined features, documenting polymorphism of these genes also presents important methodological challenges that have so far largely prevented the comprehensive analysis of complete allelic series in natural populations, and also precluded the obtention of complete genic sequences for many S-alleles. Here, we present a novel methodological approach based on a computationally optimized comparison of short Illumina sequencing reads from genomic DNA to a database of known nucleotide sequences of the extracellular domain of *SRK (eSRK)*. By examining mapping patterns along the reference sequences, we obtain highly reliable predictions of S-genotypes from individuals collected in natural populations of *Arabidopsis halleri*. Furthermore, using a *de novo* assembly approach of the filtered short reads, we obtain full length sequences of eSRK even when the initial sequence in the database was only partial, and we discover new *SRK* alleles that were not initially present in the database. When including those new alleles in the reference database, we were able to resolve the complete diploid SI genotypes of all individuals. Beyond the specific case of Brassicaceae S-alleles, our approach can be readily applied to other polymorphic loci, given reference allelic sequences are available.

## Introduction

Highly polymorphic loci across the genome play pivotal roles in important biological features (Prugnolle et al. 2005). Yet, these loci are rarely characterized comprehensively because of the technical challenges of obtaining comprehensive allelic series and accurately typing their many divergent alleles. One extreme example of this class of loci is the classical self-incompatibility (SI) system in the flowering plants, which is a common genetic mechanism to prevent self-fertilization in hermaphrodite species thanks to recognition and rejection of self-pollen by the pistils (Takayama and Isogai 2005). The genes controlling SI share some evolutionary properties with self-recognition systems in animals, such as the major histocompatibility complex (Edwards and Hedrick 1998). In many self-incompatible species, SI is controlled by a single genetic locus, the multiallelic SI locus (S-locus) that can be considered the most variable locus in plant genomes due to its extremely high allelic diversity (typically of the order of 10-35 alleles in intra-population samples) together with extreme sequence divergence among alleles of its pollen and pistil functional components, leading to high degrees of trans-specific and even trans-generic polymorphisms (Castric and Vekemans 2004). Such high allelic and sequence diversity is due to the effect of strong negative frequency-dependent selection acting on the S-locus (Wright 1939; Vekemans and Slatkin 1994). Because of such diversity, genotyping the S-locus in natural populations has been a challenge since the discovery of the pistil S-locus gene in Solanaceae (McClure et al. 1989) and in Brassicaceae (Stein et al. 1991). Although initial exploration of allelic diversity in S-locus genes involved cloning and sequencing approaches (Schierup et al. 2001), other approaches were attempted to perform systematic surveys in natural populations or in germplasm collections of domesticated species. Such approaches typically involved PCR or RT-PCR amplification using sets of general degenerate primers designed in conserved regions of the genes (Richman et al. 1995; Nishio et al. 1996; Ishimizu et al. 1999; Charlesworth 2000) followed by separation of alleles using restriction enzyme profiling (Nishio et al. 1996; Ishimizu et al. 1999; Charlesworth 2000), intron length variation (Sonneveld et al. 2006), or SSCP (Joly and Schoen 2011). One of the main problems with these approaches are that different sets of primers need to be used in order to cover the range of S-haplotype sequence divergence occurring in self-incompatible species. In addition, they often lead to incomplete genotype information because of missing (highly diverged) alleles. Alternative methods have been used in a few model species, involving systematic presence/absence PCR screening with S-allele specific primers, followed by gel electrophoresis (Broothaerts 2003; Sonneveld et al. 2003; Bechsgaard et al. 2004). Such approaches have allowed some limited surveys in natural populations in order to test for the effect of negative-frequency dependent selection on allele frequency changes across generations (Stoeckel et al. 2011), to estimate population genetic structure at the S-locus (Schierup et al. 2008), or to test for the effect of S-allele dominance on allelic frequency in sporophytic SI systems (Llaurens et al. 2008). Taking advantage of next-generation sequencing (NGS) technologies in the study of SI systems has been rather difficult because (1) amplicon-based approaches were of limited use because of the lack of universal primers, and because the size of the amplicons is constrained by the distances between conserved regions where primers are designed within the S-locus genes, which were usually larger than individual reads obtained by the dominant technologies (i.e. amplicon sizes ranging 714-1,048 bp for *Arabidopsis lyrata*, Charlesworth 2000; 318-2,500 bp for *Malus domestica*, Dreesen et al. 2010; 230-458 bp for *Prunus avium*, Sonneveld et al. 2006; and around 1,300 bp in *Brassica rapa*, Nishio et al. 1996), and (2) shotgun approaches were of limited use because alignment on a single reference sequence was hampered by high sequence divergence. Finally, the high content of repeat sequences of the S-locus region in most species where it was investigated causes important challenges for approaches relying on de novo assembly of this particular genomic region (Vekemans et al. 2014). Despite these difficulties, Jørgensen et al. 2012 proposed a method to genotype large individual samples of *Arabidopsis lyrata* using a barcoded amplicon-based method with 454 sequencing. Their method bypasses the lack of generality of the primers by using separately four pairs of general primers for each individual, followed by pooling of all PCR products, and bypasses the limit of amplicon size by sequencing only one side of the amplicons. The method has been used successfully by Mable et al. (2017, 2018) to genotype the S-locus in diploid as well as tetraploid populations of the species *A. lyrata* and *A. arenosa*. However the approach is sensible to PCR amplification biases, and the resulting coverage can be too low for some alleles to resolve reliably the sequencing errors. As an alternative approach, Tsuchimatsu et al. (2017) investigated S-locus diversity species-wide in *A. thaliana*, using individual shotgun sequencing data from 1,083 genomes, and separate mapping of short reads to each of the three available S-locus reference sequences. This approach successfully identified a diversity of recombinant S-haplotypes (Tsuchimatsu et al. 2017), but it is unclear whether its success was contingent on the low allelic diversity (only three divergent S-alleles) in the selfing *A. thaliana* and how it can be extended to species with functional SI, i.e. characterized by a large number of highly divergent S-locus allelic lines. A similar approach has been used to identify the S-locus genotype of individuals from *Malus domestica*, based on individual shotgun sequencing data and a database of 35 previously sequenced S-RNase alleles (De Franceschi et al. 2018). The approach was successful but involves manual fine-tuning and requires the availability of an exhaustive database of reference sequences, which in many species will not be available.

Here, we present a novel methodological framework building upon the approach of Tsuchimatsu et al. (2017), using data from individual shotgun sequencing in order to genotype individuals at the S-locus based on the availability of a large database of known S-allele reference sequences. The method uses two complementary approaches (see Materials and Methods for detailed information): (1) a mapping approach which attempts to map sequence reads successively on each of the reference sequences and reports a set of custom statistics to identify the best hits based on quantitative thresholds (Fig. 1A); and (2) a *de novo* assembly approach for the discovery of novel S-allele sequences and phylogenetic reconstruction to place the obtained assembled contigs within the phylogeny of S-allele reference sequences (Fig. 1B). We produced optimized pipelines to perform both analyses based on a dataset of FASTQ read sequences and a set of reference sequences. We applied the method to a published dataset consisting of 56 *A. halleri* subsp. *gemmifera* individuals for which Kubota et al. (2015) obtained genomic Illumina resequencing reads. We show that the method allows exhaustive genotyping of heterozygous as well as homozygous individuals, providing the first accurate evaluation of the frequency distribution of S-alleles in a natural population of self-incompatible Brassicaceae. Although the method has been designed and tested for studying S-allele diversity in Brassicaceae, it can be applied to any type of highly polymorphic locus, given a sample of reference allelic sequences is available.

**Fig. 1.**
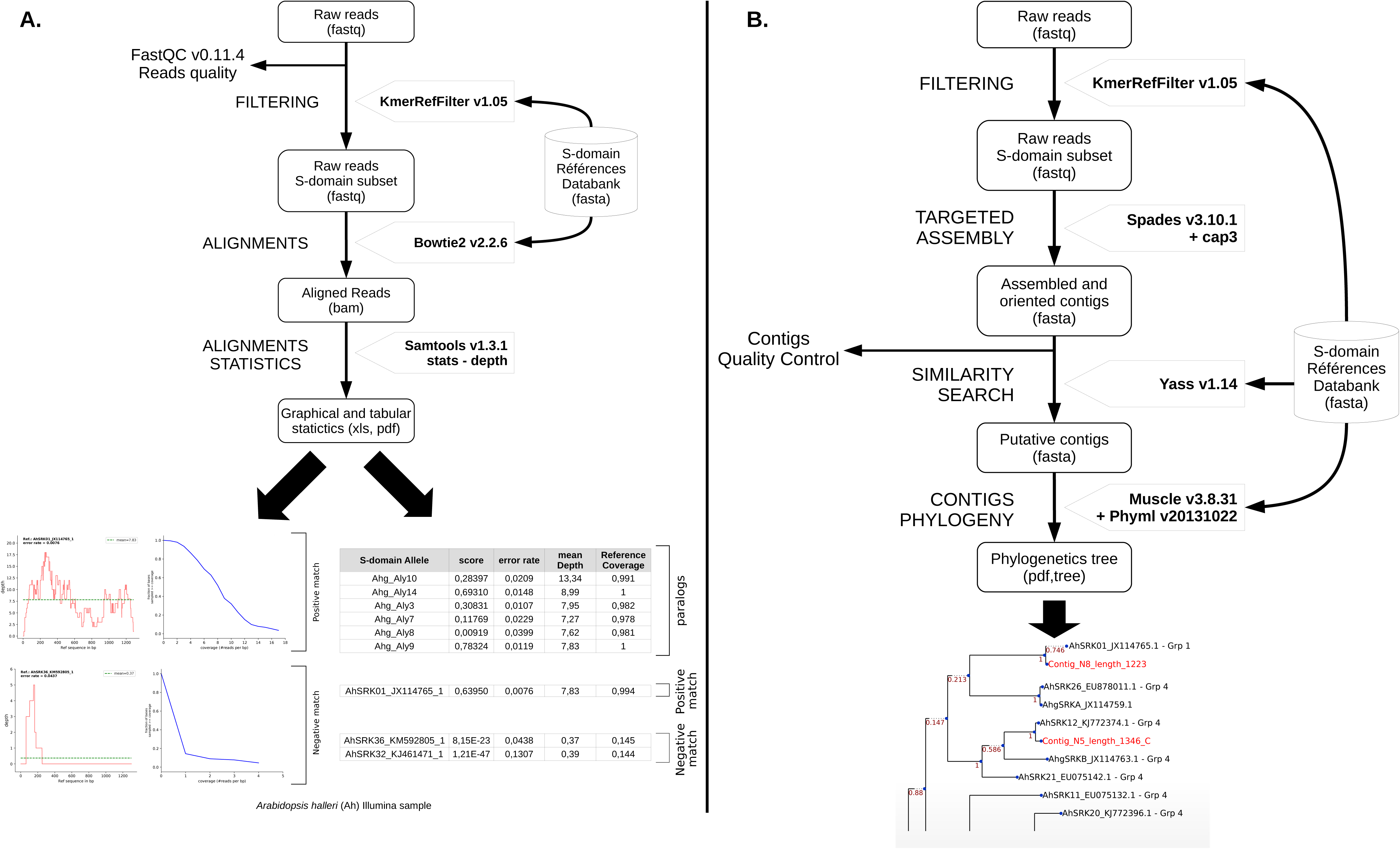
Workflow of the pipeline. **A.** Mapping approach using reference sequences and output of reporting statistics for the genotyping procedure. **B.** de *novo* assembly approach to obtain new allelic sequences.

## Material and Methods

### Database structure, curation and transformation into a k-mer library

The first step of our approach was to construct a database of publicly available nucleotide sequence variants of the gene of interest. In the present case, we started from divergent *SRK* alleles in *A. halleri* based on published studies (Castric and Vekemans 2007; Castric et al. 2008; Goubet et al. 2012). To compare mapping properties in a controlled manner and establish thresholds for genotyping, we also retrieved single-copy low diversity paralogous sequences (*Aly3, Aly7, Aly8, Aly9, Aly10, Aly14, AlLal2*) from the draft *A. halleri* subspecies *gemmifera* genome (Briskine et al. 2017). All reference sequences in the database were trimmed to include only the extracellular domain using AhSRK03 (KJ772380.1) as a reference. We then developed a dedicated algorithm (*kmerRefFilter*) to produce an exhaustive library of *k*-mers present in the set of reference sequences (size 20-mers), from which duplicates and low-complexity k-mers (using a Shannon Entropy threshold of 0.8) were removed.

### Short reads preparation and filtering

We then retrieved raw FASTQ Illumina sequence data published by Kubota et al. (2015) on 56 *A. halleri* subsp. *gemmifera* individuals from natural populations in Japan and checked their quality with *FastQC* (v0.11.4). To reduce the dataset and decrease computation time, we filtered these raw reads using the k-mer library obtained above. Paired reads were kept only if at least one k-mer matches exactly on either the forward or the reverse read sequence. This results in a drastic reduction of the number of reads (0.03% on average), allowing very efficient processing of downstream steps.

### Short reads alignment and procedure for genotype calling

The pipeline then proceeds with unpaired alignment with end-to-end sensitive mode of the reduced set of paired reads onto each sequence of the reference database using *Bowtie2 (v2.2.6)* (Langmead and Salzberg 2012). The resulting alignment files (in bam format) were analyzed using samtools (v1.4) (Li et al. 2009) to extract alignment metrics, specifically the error rate (number of mismatches divided by the number of mapped bases) reflecting the mean identity between the aligned reads and the reference sequence, and the coverage defined as the percentage of positions of the reference sequence covered by at least one read. Based on these two statistics, we aimed at identifying discriminatory criteria to distinguish positive from negative calls. To achieve this, we observed the distributions of these statistics on the paralogous sequences, that by definition will be present in two copies in diploid individuals. We fitted simple probability density functions on these empirical distributions and integrated them to obtain cumulative distribution functions. For each individual, we then computed a genotyping score against each S-allele sequence in the reference database as the product of the probability of a positive call obtained from the distribution function of the error rate times the square root probability obtained from the distribution function of the coverage. We determined the threshold for positive calls based on the empirical distribution of genotyping scores from the paralogous sequences. S-locus genotype calls can either be automated based on this threshold, or determined visually by examination of detailed mapping patterns across the reference database.

Because individuals in which a single S-allele was called can correspond to either true homozygotes or to incomplete heterozygous genotypes, we then set to determine whether S-locus homozygous calls can be distinguished from heterozygous calls. We reasoned that the depth of mapped reads for homozygous genotypes is expected to be twice that of heterozygous genotypes, and comparable in magnitude to that observed for paralogous sequences. Thus, we compared read depth, normalized for each individual by the median read depth obtained from its paralogous sequences (computed as the median of the number of reads aligned per position of the reference sequence), between individuals for which a single S-allele had been called (putative homozygotes, for which normalized read depth is expected to be close to 1), and individuals for which two distinct S-alleles had been called (putative heterozygotes, for which normalized read depth is expected to be close to 0.5).

### Allele discovery by *de novo* assembly

The genotyping procedure outlined above is based on alignment to a database of reference S-allele sequences. By definition, it is thus limited by how complete the database was to begin with. To overcome this limitation, we sought to develop a procedure to discover new S-alleles. We reasoned that *de novo* assembly of the filtered set of paired reads should potentially identify unknown S-alleles. To explore this possibility, we used the *dipSPAdes* assembler v3.10.1 (Safonova et al. 2015) using the following parameters (-k 21,41,81 and –careful) to obtain a set of assembled contigs. Then the *cap3* sequence assembly program v10/15/07 (Huang and Madan 1999) was used with default parameters to generate consensus sequences from overlaps between forward or reverse contigs. These consensus contigs were aligned against sequences of the *SRK* database using the local DNA alignment tool *YASS v1.14* (Noé and Kucherov 2005) that allows alignment even in cases of distant similarity, as appropriate given the important divergence among *SRK* alleles. We retained all contigs aligned over at least 10% of a reference and not aligning against any paralog. Query and subject alignment positions were used for contigs orientation.

As an internal validation of the assembled contigs, we examined how the paired reads mapped on them using *Bowtie2 v2.2.6* (paired alignment with end-to-end sensitive mode) (Langmead and Salzberg 2012). *Samtools v1.4* (Li et al. 2009) was used to extract several metrics, including coverage, mean read depth and depth across successive intervals (for detection of chimerism when depth levels vary sharply along the contig), insert average size, error rate (to detect cases where different S-alleles were assembled into a single hybrid contig), pairs with other orientation (used to evaluate contigs mis-assembly such as inversions), pairs on different contigs (to evaluate redundancy by detecting if paired reads align to multiple contigs). These metrics can be used to curate the set of contigs visually and evaluate assembly quality.

Finally, to compare the set of *de novo* contigs to *SRK* sequences in the starting reference database, we constructed a phylogenetic tree. Because of the procedure used for the initial filtering step of paired reads, the obtained contigs can be larger than the *SRK* sequences in the starting reference database and extend into the divergent 5’UTR and first intron. Hence, 5’ and 3’ ends were trimmed based on the largest pairwise *YASS* alignment. *MUSCLE v3.8.31* (Edgar 2004a; Edgar 2004b) was used for multiple alignment and PHYML (Guindon et al. 2010) was used for building a neighbor-joining phylogenetic tree. The program outputs one tree per individual sample, as well as one global tree where all contigs in a collection can be compared to all others and to the *SRK* sequences in the database. Once validated, the newly discovered *SRK* alleles can then be integrated in the database used for genotyping, in an iterative procedure.

## Results and Discussion

### A powerful genotyping procedure

We started from the heterogeneous database of all *SRK* sequences of *A. halleri* obtained from several published studies and deposited in NCBI. Collectively, this database contains 34 *SRK* allelic lines (Table S1), but at this step it is unknown how much of the overall diversity present in the species this represents. Because they were obtained by different molecular methods, some of these sequences cover the complete extracellular domain (n=16, average length of the complete extracellular domain = 1300 bp), but the majority of them (n=18) are partial sequences (average length = 570 bp). We mapped the raw filtered Illumina reads of the 56 *A. halleri* individuals of Kubota et al. (2015) onto the paralogs of the reference database and observed a good fit of the empirical distribution of error rates with a normal distribution and of the distribution of coverage with an exponential distribution (Fig. S1, Fig. S2A). We then mapped the filtered reads onto the S-alleles reference sequences and examined the mapping patterns (Fig. S2B). A genotyping score of 10^-5^ provided good discrimination power among S-alleles (Fig. S2B), as no individual had scores above this threshold for more than two of the reference sequences in the database, as expected for diploid genotypes (Fig. 2). Strikingly, even when the mean sequencing depth was as low as e.g. X=5, our approach allowed proper discrimination (Fig. S3). Overall, at this stage we obtained 28 full S-locus genotypes (12 homozygotes and 16 heterozygotes), 24 potentially incomplete genotypes (a single allele identified, corresponding either to homozygous individuals or heterozygous individuals in which just a single allele could be determined), while no allele could be identified for the four remaining individuals (but see below). Among the 34 *SRK* alleles present in the reference database, ten alleles were found within the analyzed dataset (AhSRK01, AhSRK02, AhSRK04, AhSRK12, AhSRK18, AhSRK19, AhSRK21, AhSRK26, AhSRK28, AhSRK-B). The large number of incomplete or missing genotypes strongly suggests that several S-alleles were absent from our reference dataset. This may partly be due to the strong divergence of the *A. halleri* subsp. *gemmifera* populations from the European *A. halleri* populations (Šrámková-Fuxová et al. 2017), as the latter were mostly used in previous screens of S-allele diversity (Castric and Vekemans 2007). Hence, we used a *de novo* assembly procedure to attempt identifying these unknown alleles.

**Fig. 2.**
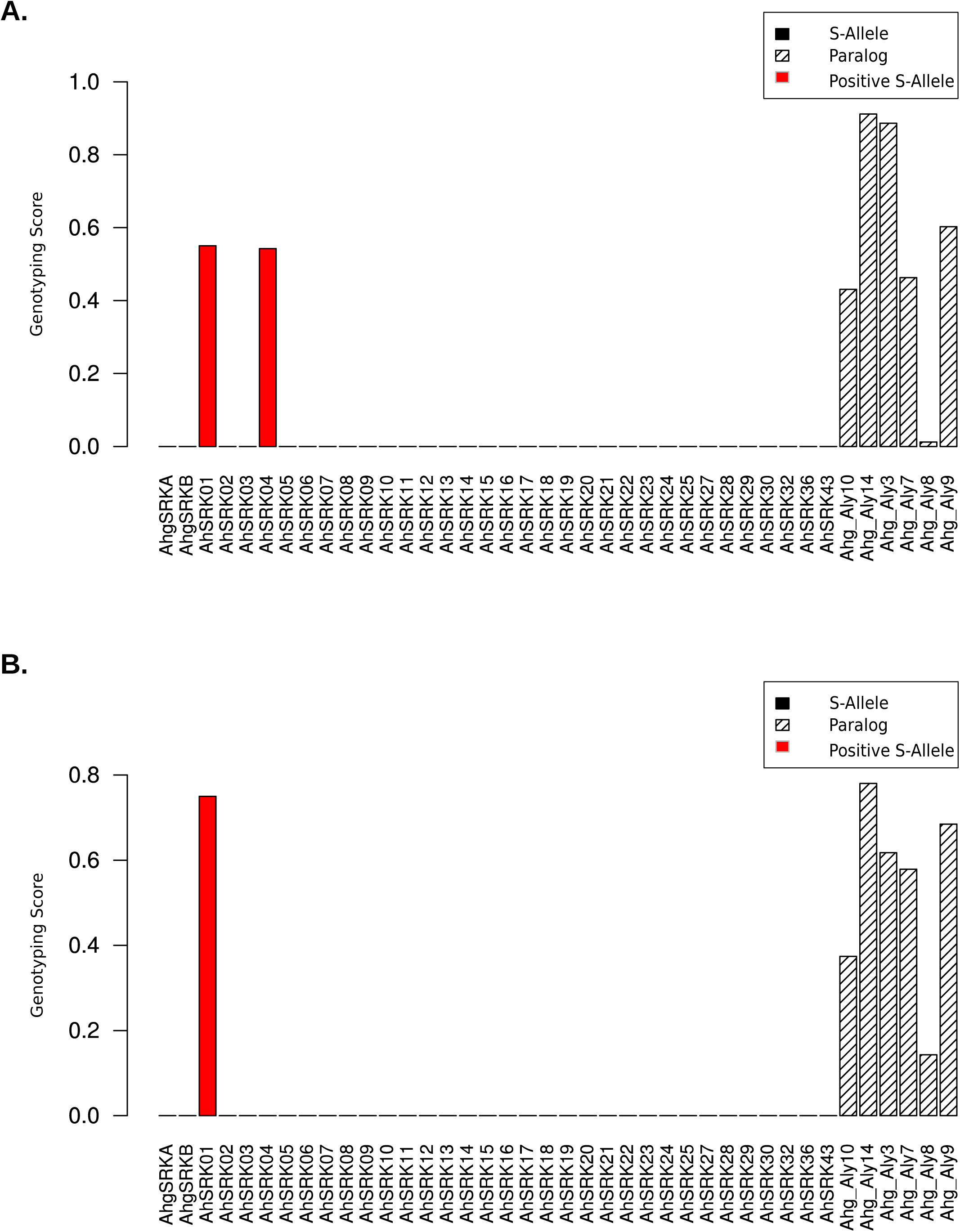
Genotyping scores for two *A. halleri* individuals for all sequences of the reference database. **A.** Example of an individual (DRS032531) with two different positive *SRK* alleles (AhSRK01/AhSRK04). **B.** Example of an individual (DRS032528) with a single positive *SRK* allele (AhSRK01)

### De novo assembly

The *de novo* assembly procedure allowed us to obtain full-length sequences of the S-domain for several *SRK* alleles that were previously only partially sequenced (Castric and Vekemans 2007). For instance, the reference sequence of AhSRK02 in GENBANK (EU075125.1) was only 550 bp long, but we obtained 4 assembled contigs with high similarity to AhSRK02, two of which appear to cover the full-length S-domain, i.e. 1294 bp. (Fig. 3A). The accuracy of the assembly is demonstrated by the strict identity of the two full-length sequences obtained (DRS032559 and DRS032547 in Fig. 3A). Overall, among the ten identified *SRK* alleles, four were present in the database as partial sequences (AhSRK02, AhSRK18, AhSRK19, AhSRK26) and we were able to obtain full-length sequences of the S-domain for all of them. Strikingly, the procedure also allowed us to recover sequences from alleles that were initially absent from the database, as illustrated in Fig. 3B. For instance, we obtained sequences of the same new allele (that we named temporarily AhNEW1) from three different individuals, which is close to but distinct from the previously known AhSRK22 allele (Fig. S4). By comparing those sequences with *SRK* sequences from the closely related species *A. lyrata*, we found that AhNEW1 is very similar to AlSRK09, which confirms that AhNEW1 and AhSRK22 are indeed distinct functional alleles, as trans-specific polymorphisms are very common at the S-locus (Fig. S4; Castric et al. 2008). Overall, the *de novo* approach identified nine new *SRK* alleles (Table S2): (1) five new alleles in *A. halleri* for which we found a trans-specific allele in *A. lyrata* (AhNEW1 - AlSRK09; AhNEW4 - AlSRK50; AhNEW5 - AlSRK23; AhNEW8 - AlSRK29; AhNEW15 - AlSRK10); and (2) four new alleles that are also unknown in *A. lyrata* (AhNEW2, AhNEW3, AhNEW7, AhNEW14) and putatively represent novel allelic lineages. The fact that at least two independent copies were observed for each *de novo* assembled allele (with the exception of AhNEW2, which was observed only once) provides high confidence in these reconstructed nucleotide sequences.

**Fig. 3.**
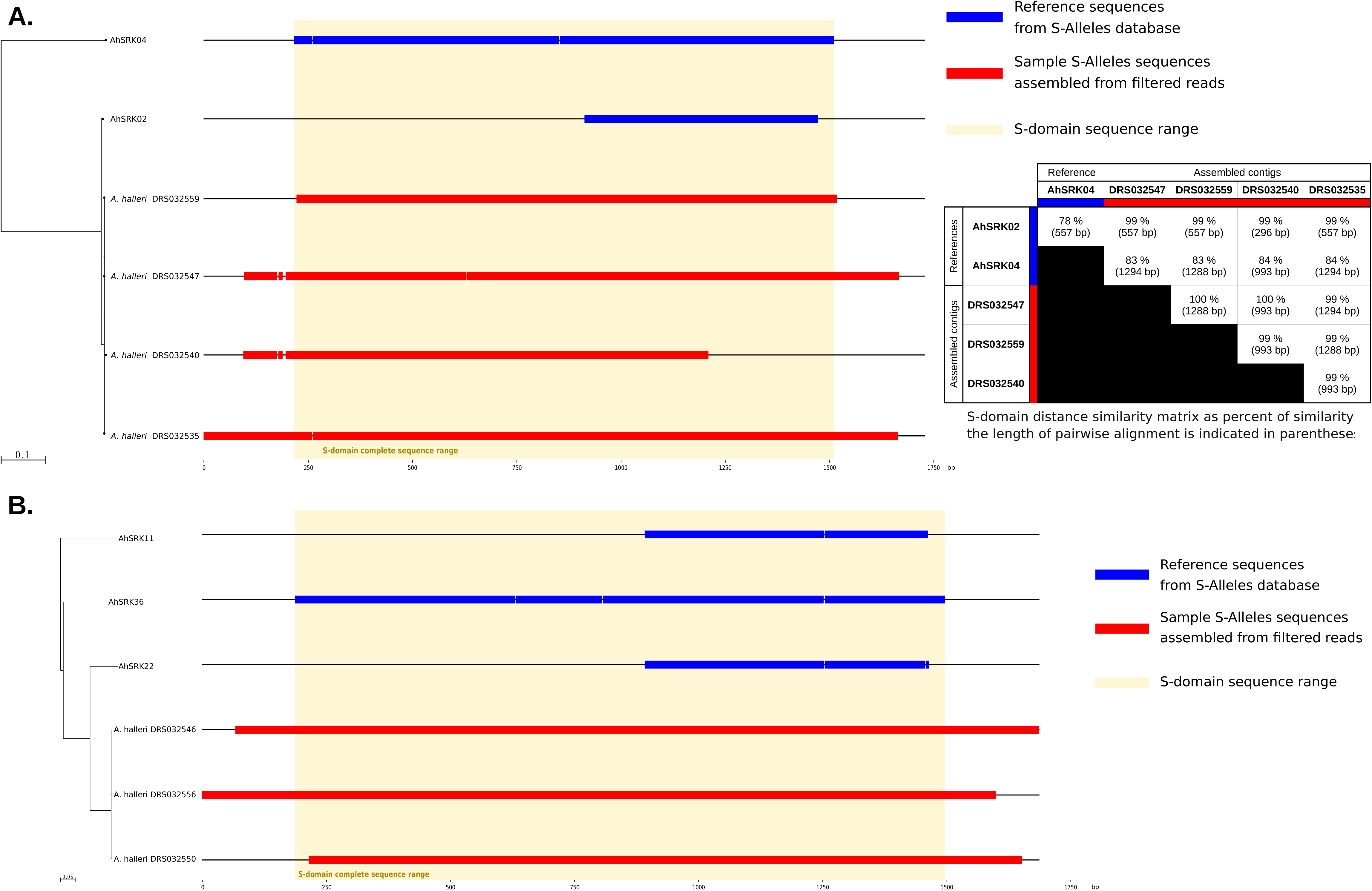
Results from the *de novo* assembly approach. **A.** S-domain full length assembly based on 4 observed copies of the AhSRK02 allele for which only an incomplete reference sequence was previously available. **B.** Discovery of a new S-allele (AhNEW1) close to but distinct from AhSRK22 and found in 3 copies in the population sample.

### A complete allele frequency distribution

Using the extended database of reference sequences that was completed by including the nine additional *SRK* alleles identified by the *de novo* assembly procedure, we compared normalized depth coverage relative to paralogs for all positive calls. We found that relative read depth for individuals with a single S-allele called was similar to that of paralogs (relative read depth close to 1, consistent with these individuals being true homozygotes), and about twice that for individuals with two S-alleles called, consistent with the latter being true heterozygotes (Fig. 2, Fig. S5, Fig. S6). Hence our approach provides a powerful way to distinguish homozygous from heterozygous individuals, and we now succeeded in obtaining full diploid *SRK* genotypes for all 56 individuals of *A. halleri* subsp. *gemmifera* (Fig. 4). The total number of *SRK* alleles observed among the 56 individuals is then 19, with 13 S-locus homozygotes (23%) and 43 heterozygotes. As expected from theory (Schierup et al. 1997; Billiard et al. 2007), homozygous genotypes were formed by alleles belonging to the most recessive classes (I and II), including the new AhNEW8 allele (Fig. 4). Overall, the total number of *SRK* alleles now described in *A. halleri* amounts to 43. In an earlier survey of *SRK* allelic diversity based on PCR screening of a large sample (N=405) from Icelandic *A. lyrata* populations, Schierup et al. (2008) found comparable estimates with 18 *SRK* alleles overall and 19.9% of homozygous individuals. As expected for sporophytic SI with dominance relationships among S-locus alleles, the allelic frequency distribution is highly skewed (Fig. S7), with the most recessive allele AhSRK01 accounting for 28.6% of allele copies, and the three alleles from the second most recessive class (AhSRK19, AhSRK28 and AhNEW8), accounting respectively for 10.7%, 10.7% and 7.1% of allele copies (Schierup et al. 1997; Billiard et al. 2007).

**Fig. 4.**
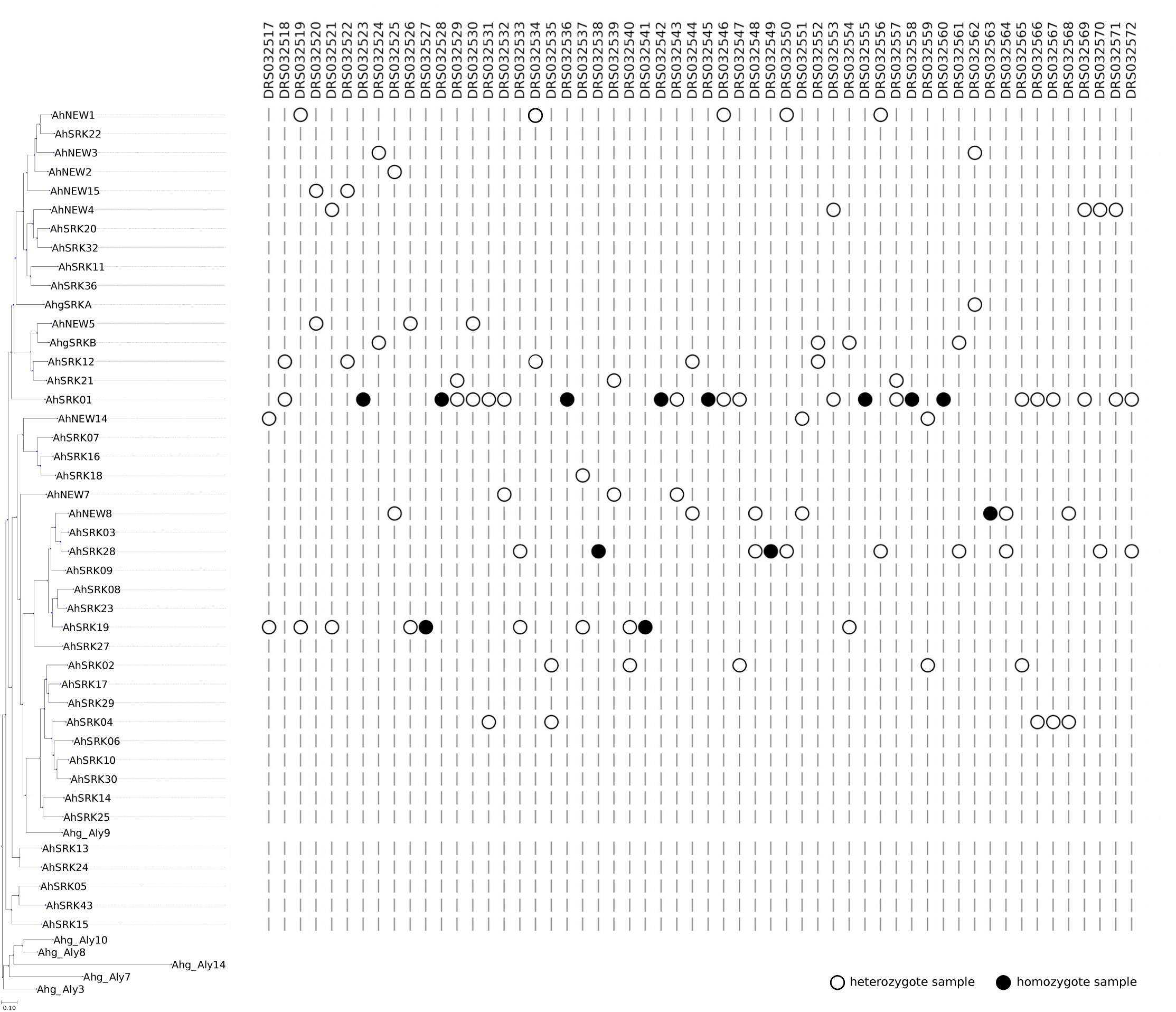
Summary of the final genotyping results on the overall sample of 56 individuals across the extended reference database including the 34 previously published SRK allelic sequences as well as the 9 new sequences. Color code: white dot (heterozygous individual); black dot (homozygous individual).

## Conclusion

We developed a powerful approach to genotype individuals at highly polymorphic loci from raw reads of shotgun individual sequence based on a sample of previously determined reference sequences. With this approach, we obtain complete sequences for new allelic lineages identified from the sampled individuals. We successfully applied this new method to the highly polymorphic SI locus of Brassicaceae and this allowed us to identify nine new S-alleles and to obtain full S-domain sequences for these new alleles as well as for four previously known alleles with partial sequences. Our approach provides an appropriate methodological framework to determine how much of the outstanding diversity of the self-incompatibility genes in outcrossing Arabidopsis remains to be discovered. This approach is readily applicable to other multiallelic genetic loci such as immune-related genes in plants or animals as an alternative to amplicon-based methods (Galan et al. 2010; Promerová et al. 2012; Grogan et al. 2016) and we provide a well-optimized ready to use script with demo data, available as an open-source project and downloadable from github: https://github.com/mathieu-genete/NGSgenotyp.

## References

Bechsgaard J, Bataillon T, Schierup MH. 2004. Uneven segregation of sporophytic self-incompatibility alleles in *Arabidopsis lyrata*: Segregation distortion in an SI-system. J. Evol. Biol. 17:554–561.

Billiard S, Castric V, Vekemans X. 2007. A general model to explore complex dominance patterns in plant sporophytic self-incompatibility systems. Genetics 175:1351–1369.

Briskine RV, Paape T, Shimizu - Inatsugi R, Nishiyama T, Akama S, Sese J, Shimizu KK. 2017. Genome assembly and annotation of *Arabidopsis halleri*, a model for heavy metal hyperaccumulation and evolutionary ecology. Mol. Ecol. Resour. 17:1025–1036.

Broothaerts W. 2003. New findings in apple S-genotype analysis resolve previous confusion and request the re-numbering of some S-alleles. Theor. Appl. Genet. 106:703–714.

Castric V, Bechsgaard J, Schierup MH, Vekemans X. 2008. Repeated adaptive introgression at a gene under multiallelic balancing selection. PLoS Genet. 4:e1000168.

Castric V, Vekemans X. 2004. Plant self-incompatibility in natural populations: a critical assessment of recent theoretical and empirical advances: self-incompatibility in natural plant populations. Mol. Ecol. 13:2873–2889.

Castric V, Vekemans X. 2007. Evolution under strong balancing selection: how many codons determine specificity at the female self-incompatibility gene *SRK* in Brassicaceae? BMC Evol. Biol. 7:132.

Charlesworth D. 2000. Population-level studies of multiallelic self-incompatibility loci, with particular reference to Brassicaceae. Ann. Bot. 85:227–239.

De Franceschi P, Bianco L, Cestaro A, Dondini L, Velasco R. 2018. Characterization of 25 full-length S-RNase alleles, including flanking regions, from a pool of resequenced apple cultivars. Plant Mol. Biol. 97:279–296.

Dreesen RSG, Vanholme BTM, Luyten K, Van Wynsberghe L, Fazio G, Roldán-Ruiz I, Keulemans J. 2010. Analysis of Malus S-RNase gene diversity based on a comparative study of old and modern apple cultivars and European wild apple. Mol. Breed. 26:693–709.

Edgar RC. 2004a. MUSCLE: multiple sequence alignment with high accuracy and high throughput. Nucleic Acids Res. 32:1792–1797.

Edgar RC. 2004b. MUSCLE: a multiple sequence alignment method with reduced time and space complexity. BMC Bioinformatics 5:113.

Edwards SV, Hedrick PW. 1998. Evolution and ecology of MHC molecules: from genomics to sexual selection. Trends Ecol. Evol. 13:305–311.

Galan M, Guivier E, Caraux G, Charbonnel N, Cosson J-F. 2010. A 454 multiplex sequencing method for rapid and reliable genotyping of highly polymorphic genes in large-scale studies. BMC Genomics 11:296.

Goubet PM, Bergès H, Bellec A, Prat E, Helmstetter N, Mangenot S, Gallina S, Holl A-C, Fobis-Loisy I, Vekemans X, et al. 2012. Contrasted patterns of molecular evolution in dominant and recessive self-incompatibility haplotypes in Arabidopsis. PLoS Genet. 8:e1002495.

Grogan KE, McGinnis GJ, Sauther ML, Cuozzo FP, Drea CM. 2016. Next-generation genotyping of hypervariable loci in many individuals of a non-model species: technical and theoretical implications. BMC Genomics 17:204.

Guindon S, Dufayard J-F, Lefort V, Anisimova M, Hordijk W, Gascuel O. 2010. New algorithms and methods to estimate maximum-likelihood phylogenies: assessing the performance of PhyML 3.0. Syst. Biol. 59:307–321.

Huang X, Madan A. 1999. CAP3: A DNA Sequence Assembly Program. Genome Res. 9:868–877.

Ishimizu T, Inoue K, Shimonaka M, Saito T, Terai O, Norioka S. 1999. PCR-based method for identifying the S-genotypes of Japanese pear cultivars. Theor. Appl. Genet. 98:961–967.

Joly S, Schoen DJ. 2011. Migration rates, frequency-dependent selection and the self-incompatibility locus in leavenworthia (brassicaceae): migration rates and frequency-dependent selection. Evolution 65:2357–2369.

Jørgensen MH, Lagesen K, Mable BK, Brysting AK. 2012. Using high-throughput sequencing to investigate the evolution of self-incompatibility genes in the Brassicaceae: strategies and challenges. Plant Ecol. Divers. 5:473–484.

Kubota S, Iwasaki T, Hanada K, Nagano AJ, Fujiyama A, Toyoda A, Sugano S, Suzuki Y, Hikosaka K, Ito M, et al. 2015. A Genome Scan for Genes Underlying Microgeographic-Scale Local Adaptation in a Wild Arabidopsis Species. PLoS Genet. [Internet] 11. Available from: https://www.ncbi.nlm.nih.gov/pmc/articles/PMC4501782/

Langmead B, Salzberg SL. 2012. Fast gapped-read alignment with Bowtie 2. Nat. Methods 9:357–359.

Li H, Handsaker B, Wysoker A, Fennell T, Ruan J, Homer N, Marth G, Abecasis G, Durbin R. 2009. The Sequence Alignment/Map format and SAMtools. Bioinformatics 25:2078–2079.

Llaurens V, Billiard S, Leducq J-B, Castric V, Klein EK, Vekemans X. 2008. Does frequency-dependent selection with complex dominance interactions accurately predict allelic frequencies at the self-incompatibility locus in *Arabidopsis halleri*? Evolution 62:2545–2557.

Mable BK, Brysting AK, Jørgensen MH, Carbonell AKZ, Kiefer C, Ruiz-Duarte P, Lagesen K, Koch MA. 2018. Adding complexity to complexity: gene family evolution in polyploids. Front. Ecol. Evol. 6:114.

Mable BK, Hagmann J, Kim S-T, Adam A, Kilbride E, Weigel D, Stift M. 2017. What causes mating system shifts in plants? *Arabidopsis lyrata* as a case study. Heredity 118:52–63.

McClure BA, Haring V, Ebert PR, Anderson MA, Simpson RJ, Sakiyama F, Clarke AE. 1989. Style self-incompatibility gene products of *Nicotlana alata* are ribonucleases. Nature 342:955–957.

Nishio T, Kusaba M, Watanabe M, Hinata K. 1996. Registration of S alleles in *Brassica campestris* L by the restriction fragment sizes of SLGs. Theor. Appl. Genet. 92:388–394.

Noé L, Kucherov G. 2005. YASS: enhancing the sensitivity of DNA similarity search. Nucleic Acids Res. 33:W540–543.

Promerová M, Babik W, Bryja J, Albrecht T, Stuglik M, Radwan J. 2012. Evaluation of two approaches to genotyping major histocompatibility complex class I in a passerine-CE-SSCP and 454 pyrosequencing: CE-SSCP vs. 454 genotyping of MHC genes. Mol. Ecol. Resour. 12:285–292.

Prugnolle F, Manica A, Charpentier M, Guégan JF, Guernier V, Balloux F. 2005. Pathogen-driven selection and worldwide HLA class I diversity. Curr. Biol. 15:1022–1027.

Richman AD, Kao T-H, Schaeffer SW, Uyenoyama MK. 1995. S-allele sequence diversity in natural populations of *Solanum carolinense* (Horsenettle). Heredity 75:405–415.

Safonova Y, Bankevich A, Pevzner PA. 2015. dipSPAdes: assembler for highly polymorphic diploid genomes. J. Comput. Biol. J. Comput. Mol. Cell Biol. 22:528–545.

Schierup MH, Bechsgaard JS, Christiansen FB. 2008. Selection at Work in Self-Incompatible *Arabidopsis lyrata*. II. Spatial Distribution of S Haplotypes in Iceland. Genetics 180:1051–1059.

Schierup MH, Mable BK, Awadalla P, Charlesworth D. 2001. Identification and characterization of a polymorphic receptor kinase gene linked to the self-incompatibility locus of *Arabidopsis* lyrata. Genetics 158:387–399.

Schierup MH, Vekemans X, Christiansen FB. 1997. Evolutionary dynamics of sporophytic self-incompatibility alleles in plants.: 12.

Sonneveld T, Robbins TP, Tobutt KR. 2006. Improved discrimination of self-incompatibility S-RNase alleles in cherry and high throughput genotyping by automated sizing of first intron polymerase chain reaction products. Plant Breed. 125:305–307.

Sonneveld T, Tobutt KR, Robbins TP. 2003. Allele-specific PCR detection of sweet cherry self-incompatibility (S) alleles S1 to S16 using consensus and allele-specific primers. Theor. Appl. Genet. 107:1059–1070.

Šrámková-Fuxová G, Záveská E, Kolář F, Lučanová M, Španiel S, Marhold K. 2017. Range-wide genetic structure of *Arabidopsis halleri* (Brassicaceae): glacial persistence in multiple refugia and origin of the Northern Hemisphere disjunction. Bot. J. Linn. Soc. 185:321–342.

Stein JC, Howlett B, Boyes DC, Nasrallah ME, Nasrallah JB. 1991. Molecular cloning of a putative receptor protein kinase gene encoded at the self-incompatibility locus of Brassica oleracea. Proc. Natl. Acad. Sci. 88:8816.

Stoeckel S, Klein EK, Oddou-Muratorio S, Musch B, Mariette S. 2011. Microevolution of s-allele frequencies in wild cherry populations: respective impacts of negative frequency dependent selection and genetic drift: selection versus genetic drift at the s-locus between two generations. Evolution 66:486–504.

Takayama S, Isogai A. 2005. Self-incompatibility in plants. Annu. Rev. Plant Biol. 56:467–489.

Tsuchimatsu T, Goubet PM, Gallina S, Holl A-C, Fobis-Loisy I, Bergès H, Marande W, Prat E, Meng D, Long Q, et al. 2017. Patterns of polymorphism at the self-incompatibility locus in 1,083 *Arabidopsis thaliana* genomes. Mol. Biol. Evol. 34:1878–1889.

Vekemans X, Poux C, Goubet PM, Castric V. 2014. The evolution of selfing from outcrossing ancestors in Brassicaceae: what have we learned from variation at the *S-* locus? J. Evol. Biol. 27:1372–1385.

Vekemans X, Slatkin M. 1994. Gene and allelic genealogies at a gametophytic self-incompatibility locus. Genetics 137:1157.

Wright S. 1939. The distribution of self-sterility alleles in populations. Genetics 24:538–552.

