## Supplemental Figures and Tables for "High throughput, high fidelity genotyping and *de novo* discovery of allelic variants at the self-incompatibility locus in natural populations of Brassicaceae from short read sequencing data"

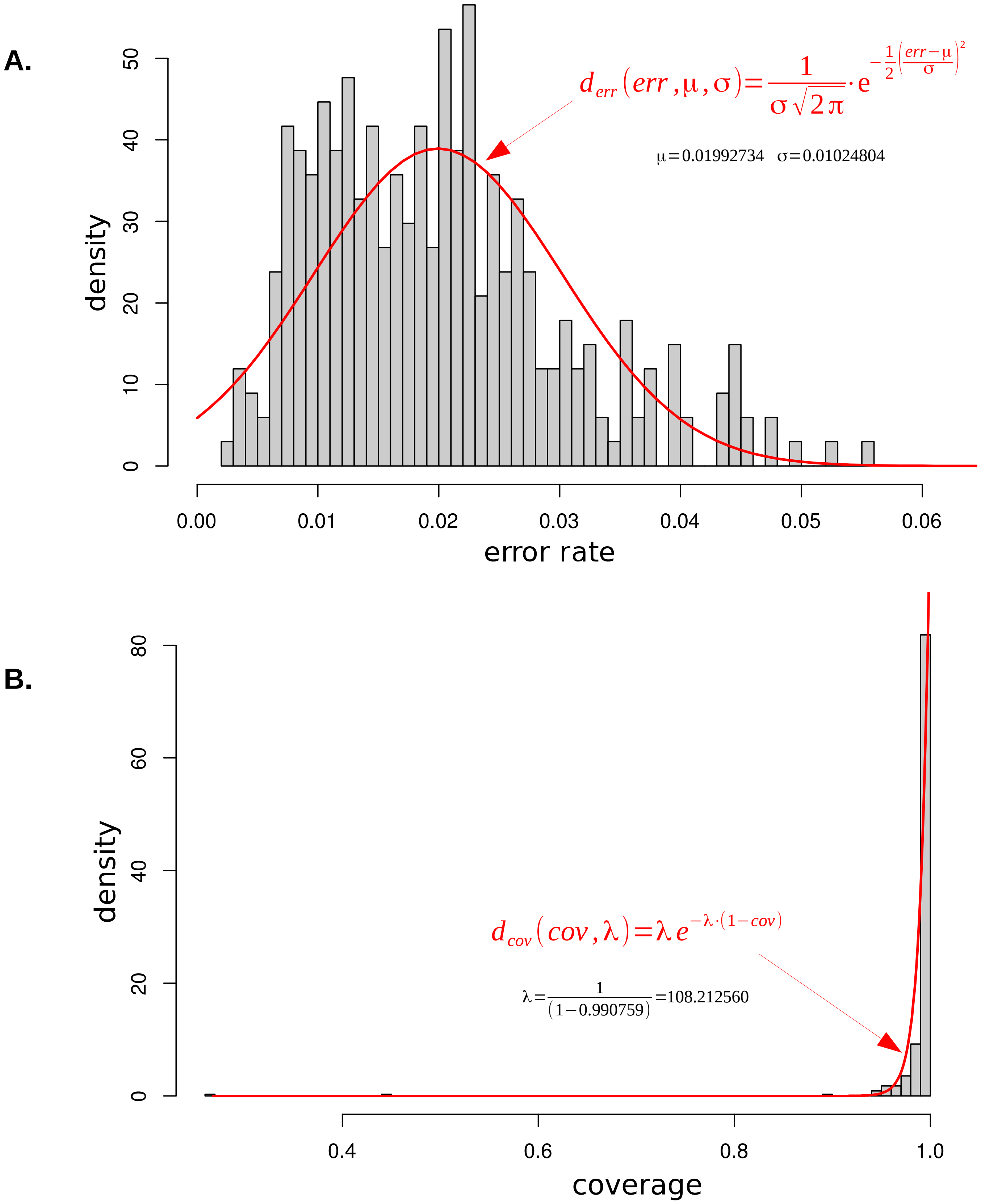


**Fig. S1.** Distributions of error rate and coverage from all alignment results between the 6 *A. halleri* subsp. *gemmifera* paralogs and the 56 samples. **A.** Error rate. The distribution of the error rate fitted to a normal function. The two parameters μ and σ, correspond respectively to the mean and standard deviation of error rate values. **B.** Coverage. The distribution of the coverage fitted to an exponential function. The parameter λ is defined as 1/(1-cov) with cov the mean of coverage values.

**A.**

**B.**

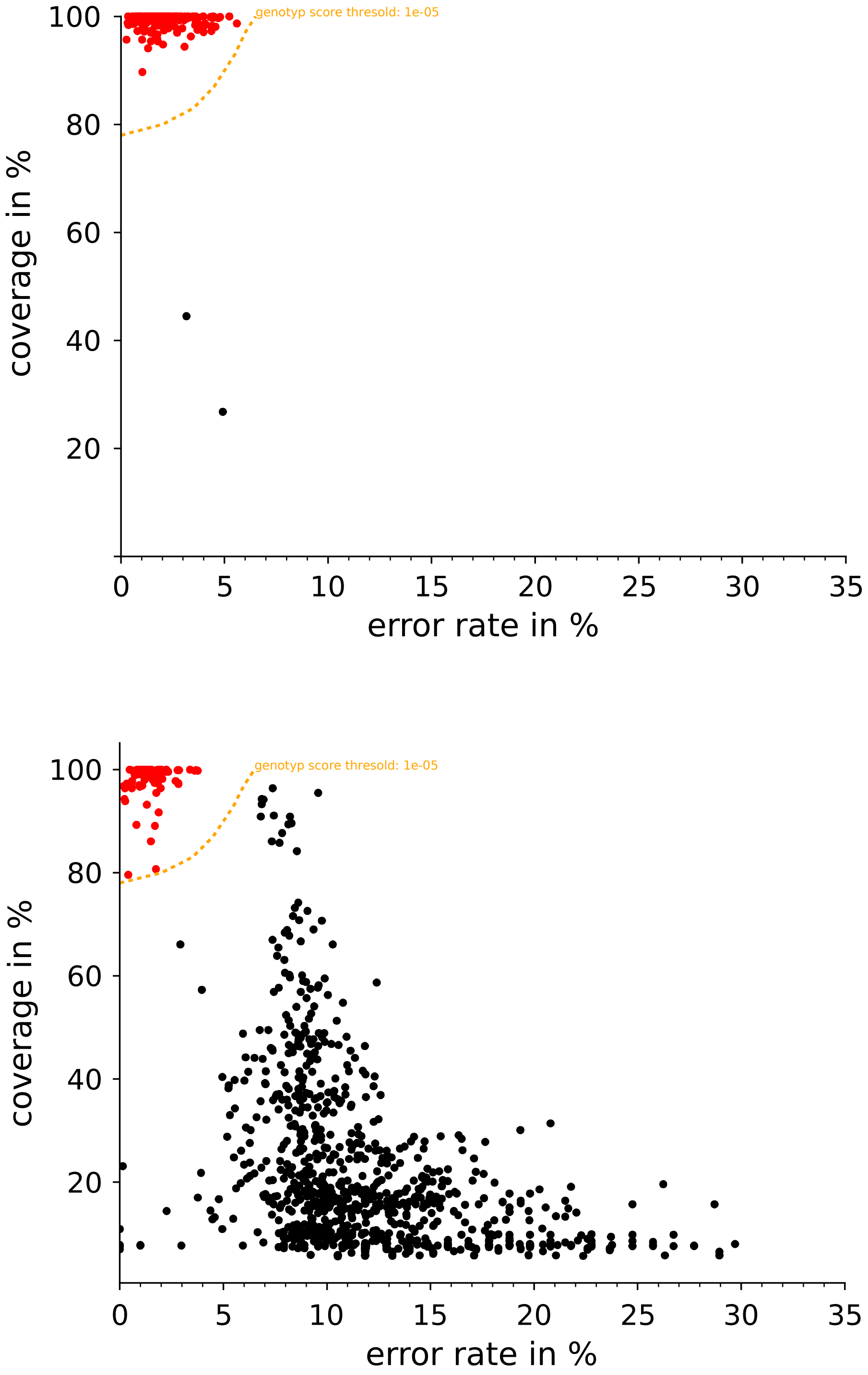


**Fig. S2.** Error rates and coverage metrics scatterplots for all alignment results between all the *A. halleri* references and the 56 samples. The genotyping score threshold (10-5) is shown as an orange dotted line. Positive (matching) and negative allele identifications are shown in red and black respectively. **A**. Paralogous sequences, for which all but two genotyping scores are high (low error rate and high coverage). The two negative points correspond to two samples that did not show positive identification for paralogue *Ahg_Aly9*. **B**. SRK references for which some references have a good match (high genotyping score, red dots) to sequencing reads of a given sample, while others have no substantial match, illustrating the good discrimination power of the genotyping score.

A.

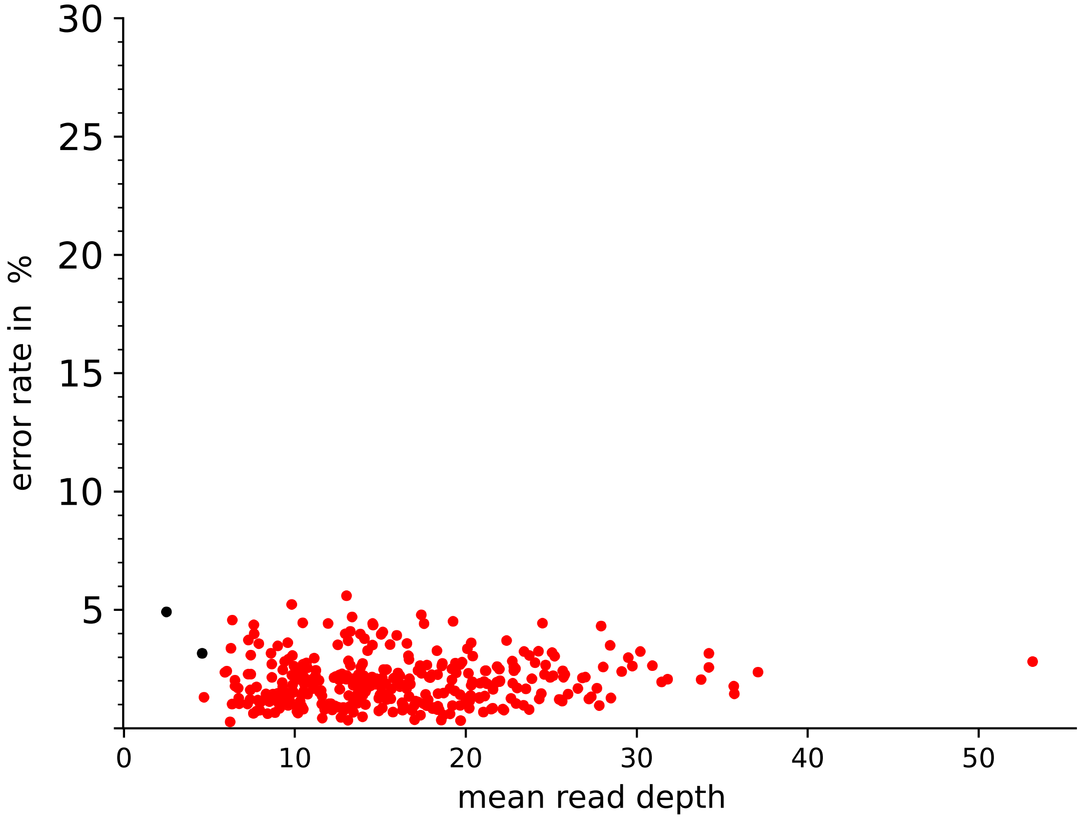


B.

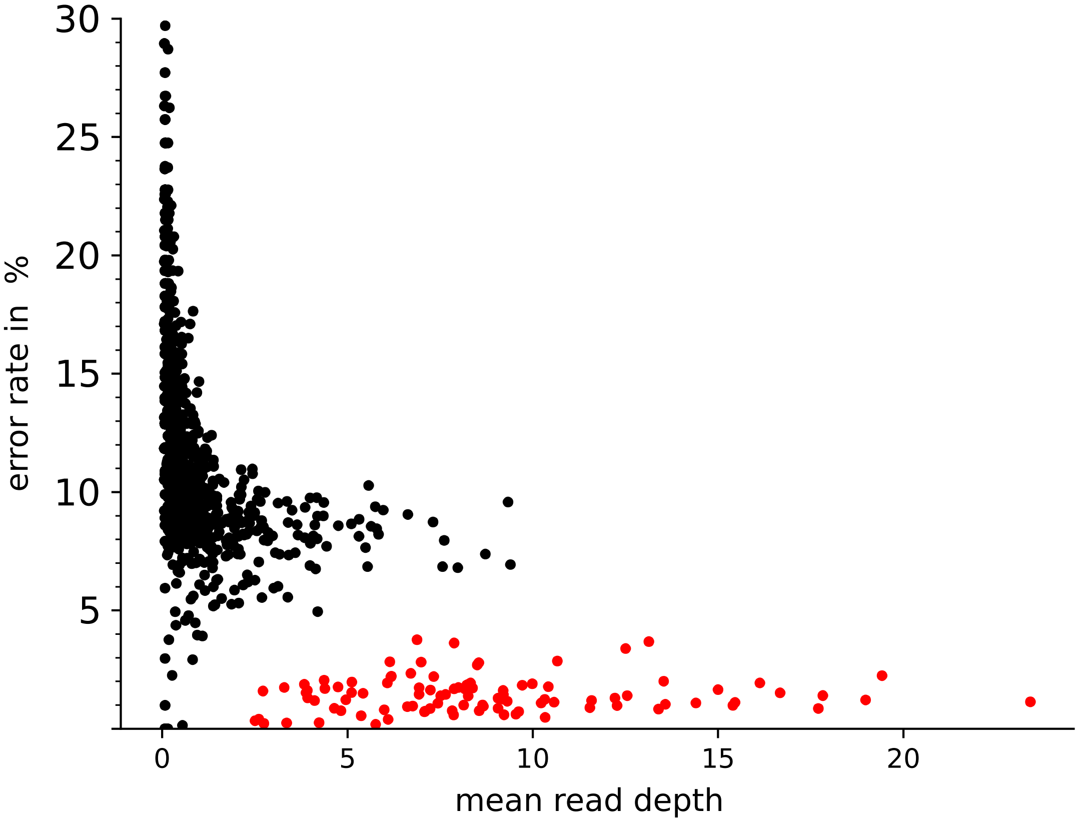


**Fig. S3.** Error rates and mean reads depth metrics scatterplots for all alignment results between all the *A. halleri* references and the 56 samples. Positive (matching) and negative allele identifications are shown in red and black respectively. **A**. Paralogous sequences. **B**. SRK references. The results show that it is possible to identify matching S-alleles even for relatively low average depth values (around 5).

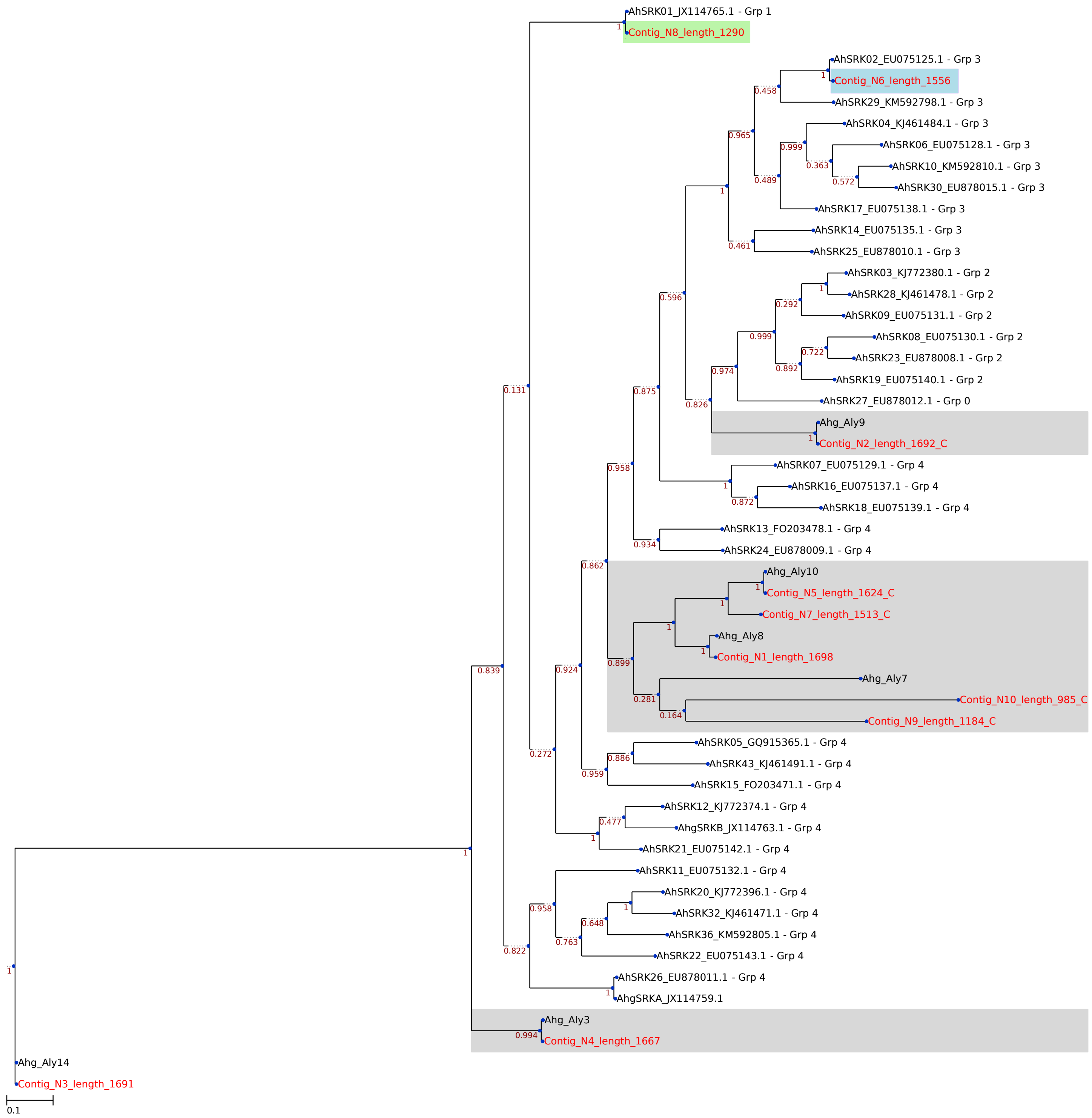


**Fig. S4.** Example of phylogeny output from the de novo assembly procedure for an S-locus heterozygous individual (DRS032547) with genotype AhSRK01 – AhSRK02. Assembled contigs are shown in red with their sequence length. Paralogs are shown in grey boxes (e.g. Contig_N4_length_1667 corresponds to the paralogous locus Ahg_Aly3). Putative S-alleles of the studied individual are shown in coloured boxes: (1) the green box corresponds to a gene copy of allele AhSRK01 from class I (Contig_N8_length_1290); (2) the blue box corresponds to a gene copy of allele AhSRK02 from class III (Contig_N6_length_1556).

**A.**

| Reads count: 6.074 | Genotyping Score | Error rate | Mean Depth | Normalized Depth | Homolog IDs | Reference Length (bp) | bases mapped | mismatches | Region Coverage |
|---|---|---|---|---|---|---|---|---|---|
| Ahg_Aly14 | 0,911436028865358 | 0,006096064 | 19,0774647887324 | 1,08129254494815 | Paralog | 1278 | 24442 | 149 | 1 |
| Ahg_Aly3 | 0,886231335469482 | 0,007560756 | 16,8466211085801 | 0,95485044863155 | Paralog | 1317 | 22220 | 168 | 1 |
| Ahg_Aly9 | 0,602641135628792 | 0,007879538 | 18,4397865853659 | 1,04514955136845 | Paralog | 1312 | 24240 | 191 | 0,993 |
| AhSRK01_JX114765_1 | 0,550119670284741 | 0,01392327 | 7,51242236024845 | 0,42579694852516 | H1001 | 1288 | 9696 | 135 | 0,995 |
| AhSRK04_KJ461484_1 | 0,542514525704374 | 0,01721107 | 8,37422360248447 | 0,47464302261186 | H3002 | 1288 | 10807 | 186 | 0,998 |
| Ahg_Aly7 | 0,462777667948495 | 0,0208849 | 16,7046632124352 | 0,94680441020361 | Paralog | 1158 | 19392 | 405 | 1 |
| Ahg_Aly10 | 0,431212332925992 | 0,02042863 | 19,185393258427 | 1,0874098278765 | Paralog | 1246 | 23937 | 489 | 0,998 |
| Ahg_Aly8 | 0,012347694596151 | 0,03701084 | 13,1157731157731 | 0,74338953568364 | Paralog | 1287 | 16968 | 628 | 0,975 |
| AhSRK10_KM592810_1 | 3,99889385787286E-18 | 0,07438436 | 3,05112316034082 | 0,17293475645379 | H3004 | 1291 | 3939 | 293 | 0,569 |
| AhSRK02_EU075125_1 | 4,36884826062604E-24 | 0,06280056 | 2,50176928520878 | 0,14179790172509 | H3001 | 1413 | 3535 | 222 | 0,212 |
| AhSRK29_KM592798_1 | 6,92034770322515E-27 | 0,08475248 | 1,94680030840401 | 0,11034278837843 | H3008 | 1297 | 2525 | 214 | 0,308 |
| AhNEW3 | 1,14916720184667E-33 | 0,08910891 | 0,21377840909091 | 0,0121167567379 | | 1408 | 303 | 27 | 0,072 |
| AhSRK30_EU878015_1 | 4,4465863874539E-34 | 0,0990099 | 0,18566176470588 | 0,01052313209756 | H3009 | 544 | 101 | 10 | 0,186 |

**B.**

| Reads count: 5.658 | Genotyping Score | Error rate | Mean Depth | Normalized Depth | Homolog IDs | Reference Length (bp) | bases mapped | mismatches | Region Coverage |
|---|---|---|---|---|---|---|---|---|---|
| Ahg_Aly14 | 0,780289519621631 | 0,01200385 | 17,8004694835681 | 1,25384031606851 | Paralog | 1278 | 22826 | 274 | 1 |
| AhSRK01_JX114765_1 | 0,750130936706858 | 0,01301092 | 12,2158385093168 | 0,86046667655046 | H1001 | 1288 | 15756 | 205 | 1 |
| Ahg_Aly9 | 0,684627016584562 | 0,01387227 | 13,969512195122 | 0,98399301221934 | Paralog | 1312 | 18382 | 255 | 0,999 |
| Ahg_Aly3 | 0,617311099411552 | 0,01593338 | 11,5368261199696 | 0,81263798811843 | Paralog | 1317 | 15251 | 243 | 0,999 |
| Ahg_Aly7 | 0,578663309589945 | 0,01789336 | 14,4240069084629 | 1,01600698778066 | Paralog | 1158 | 16766 | 300 | 1 |
| Ahg_Aly10 | 0,373933678285677 | 0,0232216 | 20,1540930979133 | 1,41962629037897 | Paralog | 1246 | 25149 | 584 | 1 |
| Ahg_Aly8 | 0,143286569430217 | 0,02821782 | 9,4048174048174 | 0,66246225911673 | Paralog | 1287 | 12120 | 342 | 0,993 |
| AhSRK19_EU075140_1 | 1,88943164259396E-22 | 0 | 0,15268329554044 | 0,01075479900772 | H2004 | 1323 | 202 | 0 | 0,076 |
| AhSRK10_KM592810_1 | 3,22585413586989E-44 | 0,1262376 | 0,31216111541441 | 0,02198819486064 | H3004 | 1291 | 404 | 51 | 0,204 |
| AhSRK02_EU075125_1 | 2,56385392507383E-45 | 0,1230552 | 0,49964614295825 | 0,03519437947339 | H3001 | 1413 | 707 | 87 | 0,098 |
| AhSRK29_KM592798_1 | 2,59221001785629E-48 | 0,1287129 | 0,07710100231303 | 0,00543088738185 | H3008 | 1297 | 101 | 13 | 0,077 |
| AhSRK04_KJ461484_1 | 3,97813658768298E-51 | 0,1386139 | 0,15683229813665 | 0,01104704898075 | H3002 | 1288 | 202 | 28 | 0,157 |

**Fig. S5.** Examples of table output from the genotyping procedure **A**. a S-locus heterozygous individual (DRS032531) with alleles AhSRK01 and AhSRK04 and **B**. an homozygote (DRS032528) with allele AhSRK01. Results for the following statistics are retrieved from Bowtie2 alignments against each sequence from the reference database: Genotyping Score (see main text), Error rate, Mean Depth, Normalized Depth, Length of the reference sequence, number of base pairs mapped on the reference over all matching reads, number of base pair mismatches, proportion of the sequence covered by at least one read. Color code: orange (matching paralogs); green (matching S-alleles, given a threshold of genotyping score of 10-5); white (non-matching S-alleles).

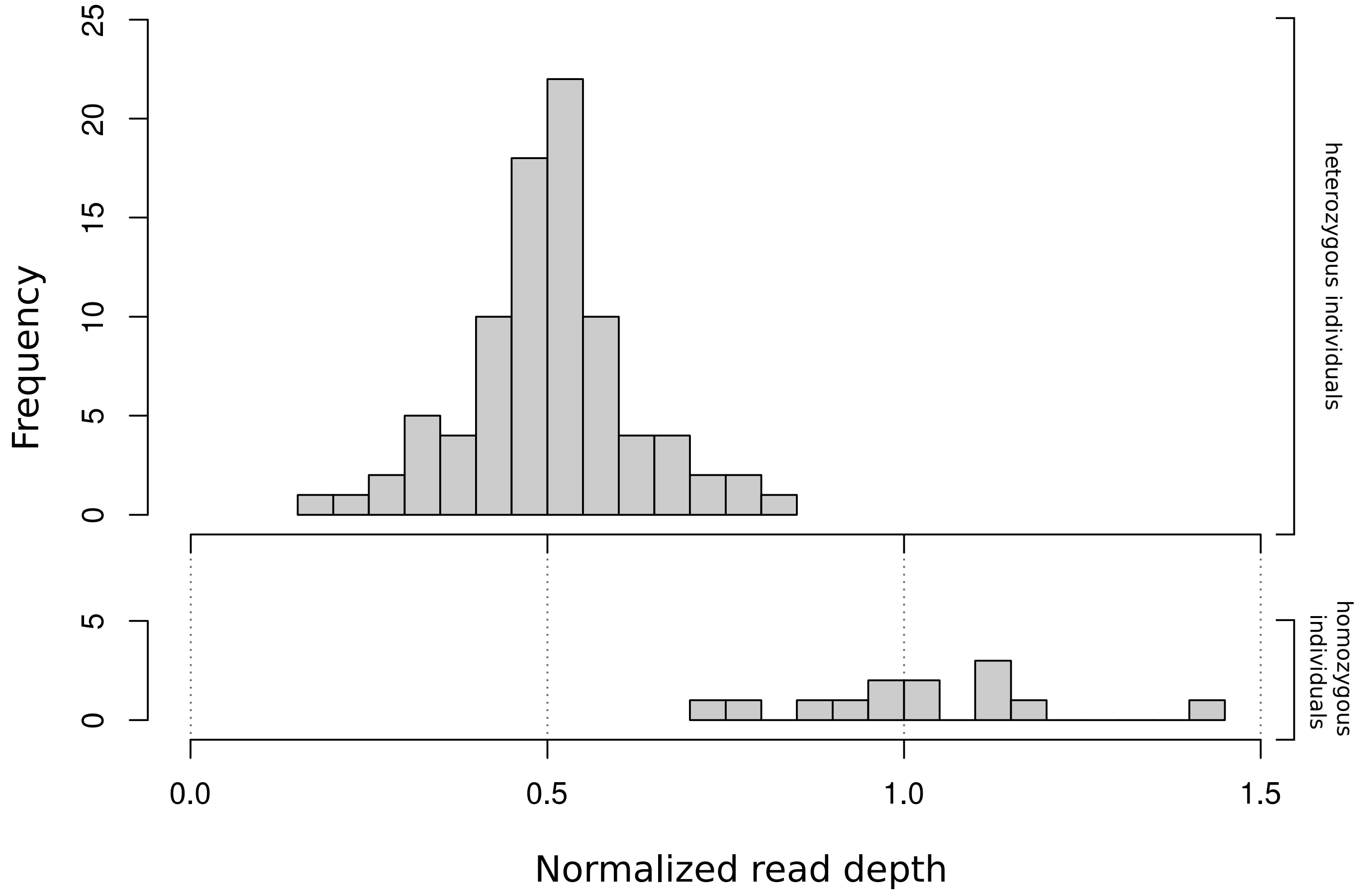


**Fig. S6.** Frequency distributions of normalized read depth against positive reference allelic sequences for individuals with two distinct S-alleles above the genotyping score threshold (top), or only one S-allele above the threshold (bottom). Normalization is performed for each individual based on the median read depth for paralogous sequences. As expected, normalized read depth is about twice as high for individuals with a single S-allele identified (putative homozygotes) than for individuals with two S-alleles identified (putative heterozygotes).

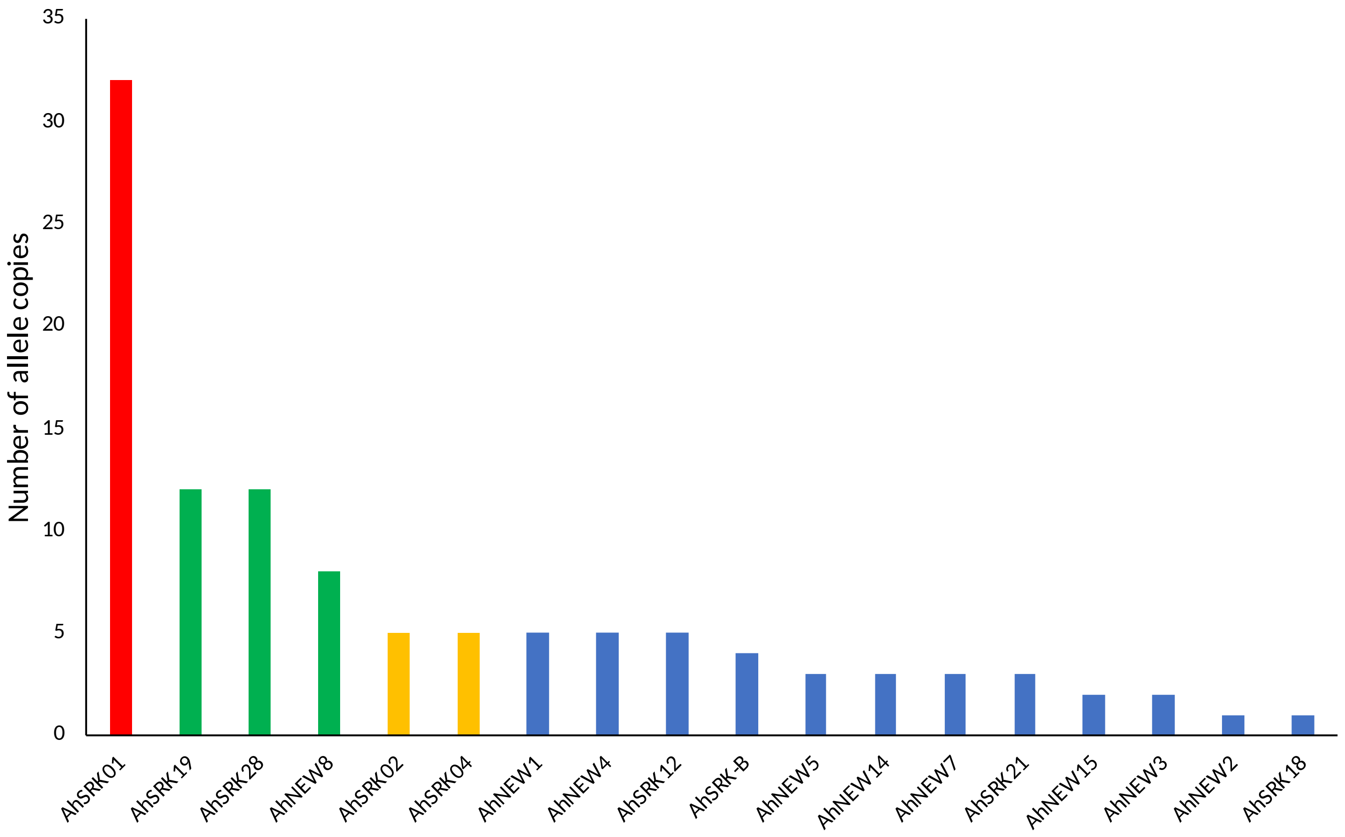


**Fig. S7.** Allelic frequency distribution across the sample of 56 individuals of *A. halleri subsp. gemmifera*. Color code: red (class I), green (class II), yellow (class III), blue (class IV). Allelic dominance increases from class I to class IV. For definition of allelic classes, see Goubet et al. (2012).

**Table S1.** List of reference S-domain sequences for the SRK alleles and list of paralogous sequences from A. lyrata used to blast the *A. halleri* subsp. *gemmifera* genome

| Allele/gene name | Sequence length | GenBank accession |
|---|---|---|
| *SRK alleles* | | |
| AhgSRKA | 1294 | JX114759.1 |
| AhgSRKB | 1300 | JX114763.1 |
| AhSRK01 | 1288 | JX114765.1 |
| AhSRK02 | 550 | EU075125.1 |
| AhSRK03 | 1321 | KJ772380.1 |
| AhSRK04 | 1288 | KJ461484.1 |
| AhSRK05 | 529 | GQ915365.1 |
| AhSRK06 | 694 | EU075128.1 |
| AhSRK07 | 607 | EU075129.1 |
| AhSRK08 | 567 | EU075130.1 |
| AhSRK09 | 573 | EU075131.1 |
| AhSRK10 | 1291 | KM592810.1 |
| AhSRK11 | 567 | EU075132.1 |
| AhSRK12 | 1291 | KJ772374.1 |
| AhSRK13 | 1285 | FO203478.1 |
| AhSRK14 | 561 | EU075135.1 |
| AhSRK15 | 1294 | FO203471.1 |
| AhSRK16 | 573 | EU075137.1 |
| AhSRK17 | 567 | EU075138.1 |
| AhSRK18 | 1300 | EU075139.1 |
| AhSRK19 | 573 | EU075140.1 |
| AhSRK20 | 1327 | KJ772396.1 |
| AhSRK21 | 570 | EU075142.1 |
| AhSRK22 | 567 | EU075143.1 |
| AhSRK23 | 558 | EU878008.1 |
| AhSRK24 | 552 | EU878009.1 |
| AhSRK25 | 546 | EU878010.1 |
| AhSRK27 | 561 | EU878012.1 |
| AhSRK28 | 1315 | KJ461478.1 |
| AhSRK29 | 1297 | KM592798.1 |
| AhSRK30 | 544 | EU878015.1 |
| AhSRK32 | 1303 | KJ461471.1 |
| AhSRK36 | 1300 | KM592805.1 |
| AhSRK43 | 1303 | KJ461491.1 |
| *Paralogous genes* | | |
| Ahg_Aly3 | 1317 | AY186752.1 * |
| Ahg_Aly7 | 1158 | AY186754.1 * |
| Ahg_Aly8 | 1287 | AY186755.1 * |
| Ahg_Aly9 | 1312 | AY186756.1 * |
| Ahg_Aly10 | 1246 | AY186761.1 * |
| Ahg_Aly14 | 1278 | AY186762.1 * |

* Genbank accession numbers for paralogous genes correspond to the *A. lyrata* sequences which have used to blast the *A. halleri* subsp. *gemmifera* genome

**Table S2.** List of new S-alleles identified in *A. halleri* with their attributed name and GenBank accession number

| **Newly identified S-Allele** | **Allele name** | **GenBank accession** | **Trans-specific allele in *A. lyrata*** |
|---|---|---|---|
| AhNEW1 | AhSRK34 | | AlSRK09 |
| AhNEW2 | AhSRK44 | | |
| AhNEW3 | AhSRK41 | | |
| AhNEW4 | AhSRK50 | | AlSRK50 |
| AhNEW5 | AhSRK42 | | AlSRK23 |
| AhNEW7 | AhSRK40 | | |
| AhNEW8 | AhSRK45 | | AlSRK29 |
| AhNEW14 | AhSRK46 | | |
| AhNEW15 | AhSRK35 | | AlSRK10 |
| AhgSRKA* | AhSRK26 | | AlSRK22 |
| AhgSRKB* | AhSRK47 | | |

* S-allele previously identified in *A. halleri* subsp. *gemmifera* but without attribution of a serial allele number in the AhSRKXX format
